# Molecular control of gene expression by *Brucella* BaaR, an IclR-family repressor

**DOI:** 10.1101/251942

**Authors:** Julien Herrou, Daniel M. Czyż, Aretha Fiebig, Jonathan W. Willett, Youngchang Kim, Ruiying Wu, Gyorgy Babnigg, Sean Crosson

**Affiliations:** Department of Biochemistry and Molecular Biology, University of Chicago, Chicago, Illinois, USA; Howard Taylor Ricketts Laboratory, University of Chicago, Argonne, Illinois, USA; Argonne National Laboratory, Argonne, Illinois, USA; Department of Microbiology, University of Chicago, Chicago, Illinois, USAx

**Keywords:** *Brucella abortus*, sigma factor, RpoE1, stress response, IclR, adipic acid, µ-aminocaproic acid, tetradecanedioic acid, µ-caprolactone, ²-oxidation, transcriptional regulation

## Abstract

The *Brucella abortus* general stress response sigma factor, σ^E1^, directly and indirectly regulates the transcription of dozens of genes that influence stress survival and host infection. Characterizing the functions of σ^E1^ regulated genes therefore contributes to understanding of *B. abortus* physiology and infection biology. Transcription of the IclR family regulator, Bab2_0215, is indirectly activated by σ^E1^ but its function remains undefined. We present a structural and functional characterization of Bab2_0215, which we have named *<u>B</u>rucella* <u>a</u>dipic acid activated regulator (BaaR). BaaR adopts a classic IclR-family fold and directly regulates the transcription of two operons with predicted roles in carboxylic acid oxidation. BaaR binds two sites on chromosome II between baaR and a divergently transcribed hydratase/dehydrogenase (*acaD2*), and represses transcription. We identified three carboxylic acids (adipic acid tetradecanedioic acid, ε-aminocaproic acid) and a lactone (ε-caprolactone) that enhance transcription from the *baaR* and *acaD2* promoters. However, neither the activating acids nor caprolactone enhance transcription by binding directly to BaaR. Induction of *baaR* transcription by adipic acid requires the gene *bab2_0213*, which encodes a major facilitator superfamily transporter, suggesting that Bab2_0213 transports adipic acid across the inner membrane. We conclude that a set of structurally related organic molecules activate transcription of genes repressed by BaaR. Our study provides molecular-level understanding of a gene expression program regulated downstream of σ^E1^.

## INTRODUCTION

Microbes use numerous mechanisms to modulate their physiology in response to changes in the intracellular and extracellular environments (1). A fundamental mechanism required for physiological adaptation is the regulation of gene expression. The Gram-negative bacterium, *Brucella abortus*, is a facultative intracellular pathogen that can cause abortion in mammals, and is among the most common zoonotic pathogens globally (2,3). *B. abortus* encodes an extracytoplasmic function-type sigma factor, σ^E1^, which directly and indirectly regulates transcription of approximately 100 genes in response to environmental perturbation. σ^E1^-dependent transcription confers resistance to multiple environmental stressors and is required for maintenance of chronic *B. abortus* infection in a mouse model (4–7). Assigning physiological and/or biochemical functions to σ^E1^-regulated genes is therefore important for understanding *B. abortus* stress physiology and infection biology.

Though several genes in the *B. abortus* σ^E1^ regulon have been characterized (4,5,7–10), the functions of the majority of σ^E1^-regulated genes remain undefined. Gene *bab2_0215*, which encodes a predicted IclR-family transcriptional regulator, was identified as having significantly reduced expression in a σ^E1^ deletion strain (*ΔrpoE1*) (4). IclR proteins are conserved across bacteria and archaea and have been reported to control the transcription of genes involved in a range of processes (11-;13), including carbon catabolism (14-;20), biofilm formation (21), quorum sensing (22,23), virulence (24-;26), stress response (27), antibiotic resistance (28,29), amino acid and secondary metabolite biosynthesis (30-;32), motility (33), and sporulation (34,35). However, the functional roles of IclR-family regulators in *Brucella* spp. remain largely unexplored. Herein, we present a structural and functional characterization of Bab2_0215, which we have named *<u>B</u>rucella* <u>a</u>dipic acid <u>a</u>ctivated <u>r</u>egulator, or BaaR. The experimental data presented in this manuscript define BaaR as a transcriptional repressor, identify the regulatory targets of BaaR, and reveal chemical signals that derepress transcription of genes inhibited by BaaR *in vivo*.

Specifically, we have solved an X-ray crystal structure of BaaR, which revealed a classic IclR fold (11) with a C-terminal ligand binding domain (LBD) and an N-terminal helix-turn-helix (HTH) DNA-binding domain. We demonstrate that deleting *baaR* (Δ*baaR*) results in strong transcriptional upregulation of four genes located adjacent to *baaR* on chromosome II of *B. abortus*. This regulated gene set shares sequence similarity with characterized dicarboxylic acid (*dca*) β-oxidation operons from *Acinetobacter* and *Pseudomonas* (36-;39). We found that BaaR functions as a transcriptional repressor by binding to two conserved palindromic motifs between *baaR* and *bab2_0216* (previously annotated as *acaD2* (<u>a</u>cyl-<u>C</u>o<u>A</u> dehydrogenase 2) (40)). A set of structurally related organic molecules enhanced transcription from a BaaR-dependent transcriptional reporter *in vivo*. Activation of transcription by one of these molecules, adipic acid, required the transporter gene *bab2_0213*, providing evidence that *bab2_0213* encodes an adipic acid transporter. However, none of the small molecules that activate transcription *in vivo* were found to bind directly to purified BaaR *in vitro*. We thus conclude that the molecular signal(s) that directly control BaaR-dependent transcription in the cell are distinct from the activating molecules we identified via selective addition to the growth medium.

## RESULTS

### Regulation of *baaR* by σ^E1^ is likely indirect, and *baaR* does not contribute to *B. abortus* hydrogen peroxide (H_2_O_2_) resistance

Previously, wild-type (WT) *B. abortus* strain 2308 and *B. abortus* 2308 harboring an in-frame deletion of the general stress regulator *rpoE1* (encoding σ^E1^) were subjected to H_2_O_2_ stress, and differences in gene expression between the two strains were assessed by RNAsequencing(RNA-seq)(4). Transcription of *baaR* was decreased two-fold in the Δ*rpoE1* strain relative to the WT strain (Figure 1A), suggesting that *baaR* is directly or indirectly activated by σ ^E1^. We could not identify a σ^E1^-binding site in the *baaR* promoter region, which suggests that this regulatory effect is indirect. To evaluate the contribution of *baaR* to σ^E1^-dependent protection against H_2_O_2_ stress, we measured the survival of the WT, Δ*rpoE1*, and Δ*baaR* strains after treatment with 5 mM H_2_O_2_. Survival of Δ*rpoE1,* assessed by enumerating colony forming units (CFU) on solid medium after hydrogen peroxide treatment, was reduced by approximately 1 order of magnitude while Δ*baaR* survival did not differ from WT survival (Figure 1B). We thus concluded that decreased expression of *baaR* in Δ*rpoE1* does not contribute to the viability defect of Δ*rpoE1* under the assayed condition.

**Figure 1:**
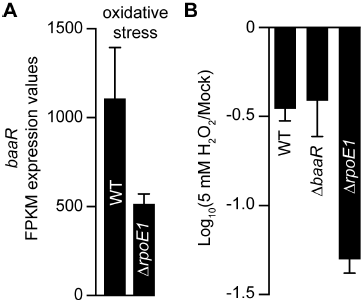
Transcriptional regulation of *bab2_0215* (*baaR*) under oxidative stress. A) The fragments per kilobase of transcript per million mapped reads (FPKM) values for *baaR* determined by RNA-seq in the *B. abortus* WT strain and a sigma factor σ ^E1^ deletion strain (Δ*rpoE1*) under oxidative stress conditions (5 mM H_2_O_2_). Error bars represent the standard deviations of three independent experiments. B) Relative survival of WT, Δ*baaR*, and Δ*rpoE1* strains after treatment with 5 mM H_2_O_2_ for 1 hour. Error bars represent the standard deviations.

### BaaR adopts a classic IclR family fold

To better understand the structure and function of BaaR, we expressed, purified, and crystallized the BaaR protein and solved its structure to 1.95 Å resolution (R_free_ = 21.2%, R_work_ = 17.7%) (see Table S1 for data collection and refinement statistics). Two BaaR dimers were present in the crystallographic asymmetric unit, and each monomer adopted the classic fold of IclR proteins (11). The monomeric protein consisted of an N-terminal winged HTH domain (residues F39-D101), α-helical linker (α4, residues I102-L115), and C-terminal LBD (residues M116-P284) (Figure 2A, B, and C). The HTH DNA-binding domain contains three α-helices (α1, α2, and α3) and two β-strands (β1 and β2) (Figure 2A and C). Remarkably, in comparison with typical IclR protein sequences, BaaR has an unusually long N-terminal region. This extension is approximately 38 amino acids (from residue M1-Q38) and is similar in size to the N-termini of *Acinetobacter* DcaR and DcaS and *Pseudomonas* P1630 (Figure 2C and Figure S1). For crystallization, a trimmed version (residues K21-P284) of BaaR was used; the N-terminus appeared to be mostly unstructured with the exception of residues G31-D36, which adopted an α-helical fold (α1’) in three out of four monomers (Figure 2A and C). The functional relevance of this region is unknown; we cannot exclude the possibility that an alternative start codon is used for protein translation *in vivo*.

**Figure 2:**
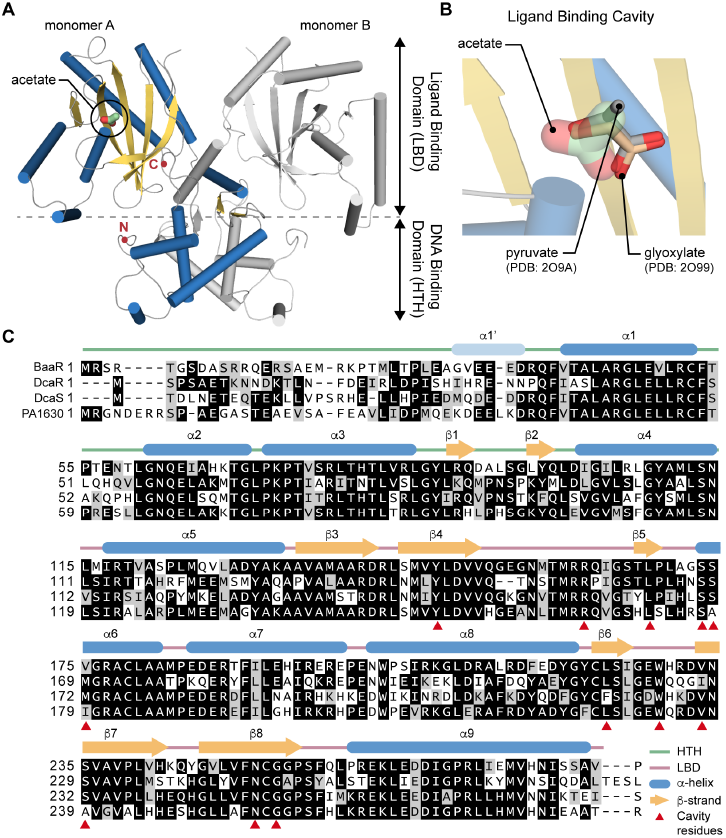
X-ray crystal structure of the *B. abortus* IclR protein, BaaR (Bab2_0215; residues 21-284). A) BaaR adopts the classic fold characteristic of dimeric IclR proteins. The dimer (monomer A + monomer B) contains an N-terminal DNA-binding domain with a winged helix-turn-helix fold (HTH; F39-D101), α-helical linker (α4, residues I102-L115), and a C-terminal ligand-binding domain (LBD; M116-P284). The K21-Q38 portion of the N-terminal extension containing α-helix α1’ is also represented. An acetate molecule, likely derived from the crystallization buffer, was found in the LBD of each monomer. In monomer A, β-strands are represented as yellow arrows and α-helices as blue cylinders. Monomer B is in gray. The C‐ and N-termini of monomer A are in red. B) Close-up view of the LBD cavity in monomer A. The acetate molecule superimposes with glyoxylate and pyruvate, two ligands found in the *E. coli* IclR LBD cavity. PDB structures 5WHM, 2O9A, and 2O99 were used for this structural alignment and correspond to the *B. abortus* BaaR structure and *E. coli* IclR structures bound to glyoxylate and pyruvate, respectively. C) Amino acid sequence alignment of *B. abortus* BaaR (Bab2_0215), *Acinetobacter* DcaR (ACIAD1688) and DcaS (ACIAD1684), and *Pseudomonas* DcaR (PA1630). Residues in the black boxes are identical, residues in the gray boxes are homologous, and residues in white do not share any homology. *B. abortus* BaaR secondary structures are shown above the alignment; β-strands are represented as yellow arrows and α-helices as blue cylinders. A light green line and light red line delimit the HTH domain and LBD, respectively. Residues found in the *B. abortus* BaaR LBD cavity are indicated by red triangles. For each amino acid sequence, the residues are numbered, starting with methionine.

The BaaR LBD consists of five α-helices (α5, α6, α7, α8, and α9) and seven anti-parallel β-strands (β3, β4, β5, β6, β7, β8, and β9) arranged in a semi-circular β-scaffold. The β-scaffold is partially occluded by α6 and folds to form a cavity in the LBD (Figure 2A and C). In the LBD cavity of each IclR protein, we observed extra electron density consistent with a bound acetate molecule (Figure 2A and B). This molecule is likely derived from the crystallization solution, which contained 200 mM calcium acetate. Notably, the acetate occupied the same position as pyruvate or glyoxylate molecules that have previously been reported to bind to the LBD of *Escherichia coli* IclR (41,42) (Figure 2B).

Protein sequence alignment of BaaR, *Acinetobacter* DcaR (ACIAD1688) and DcaS (ACIAD1684), and *Pseudomonas* PA1630 revealed high sequence homology and common secondary structural features (Figure 2C). All have HTH domains with long N-termini; these proteins also have LBD cavities comprising very similar residues, suggesting that they recognize and respond to identical or similar small molecules and interact with target DNA in a comparable manner (Figure 2C and S1). The two BaaR dimers present in the asymmetric unit are highly superimposable and displayed very few structural differences (RMSD = 0.76). IclR-family proteins have been found to adopt dimeric or tetrameric conformations in X-ray structures, in solution, or in their interactions with DNA (11,41,43-;45). Therefore, even if BaaR crystallizes as a dimer *in vitro*, it potentially exists in different oligomeric states *in vivo*.

### BaaR represses the transcription of a set of genes with similarities with the *dca* operon

Our crystal structure clearly identified BaaR as an IclR-family transcriptional regulator capable of binding a carboxylic acid (acetate) via its LBD. To identify genes specifically regulated by BaaR, we measured the global transcriptomic profile of the Δ*baaR* strain using RNA-seq and compared it with that of the WT strain (Table S2). Among the differentially expressed genes, four genes were highly upregulated in Δ*baaR* relative to WT (Figure 3A). These four genes, *bab2_0213*, *bab2_0214*, *bab2_0216*, and *bab2_0217*, are contiguous and adjacent to *baaR* (*bab2_0215*). Genes *bab2_0213*, *bab2_0214* and *bab2_0215* are located on the opposite strand from *bab2_0216* and *bab2_0217*, and the two gene sets are likely divergently transcribed (Figure3B). *bab2_0213* is annotated as a MucK transporter, a member of the major facilitator superfamily (MFS). A transporter related to *bab2_0213* in *Acinetobacter* sp. ADP1, is involved in *cis,cis* muconic acid uptake (39). *bab2_0214* is annotated as an acyl-CoA dehydrogenase; this family of flavoproteins catalyzes α,β-dehydrogenation of fatty acid acyl-CoA conjugates (46,47). *bab2_0216* and *bab2_0217* are annotated as pseudogenes. However, this annotation appears to be incorrect: a search of Pfam (48) using the primary structures of these loci suggested that *bab2_0216* (*acaD2*), is in fact two fused, in-frame genes that correspond to an enoyl-CoA hydratase/isomerase and a 3-hydroxyacyl-CoA dehydrogenase. Both of these enzymes are predicted to be involved in β-oxidation of fatty acids (49,50). Our transcriptomic data corroborate a recent proteomic study in *B. abortus* in which a small peptide corresponding to the Bab2_0216 N-terminus was identified by mass spectrometry (51), suggesting that the full-length “fusion” protein is expressed.

**Figure 3:**
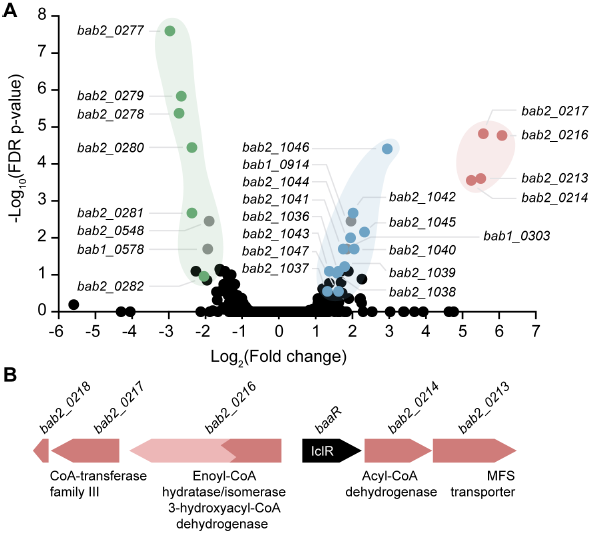
Volcano plot representing the RNA-seq transcriptomic profiles of the *B. abortus* WT strain compared with the Δ*baaR* strain. A) Gene transcription levels were compared between a *B. abortus* strain harboring a deletion in *baaR* and the WT strain. The x-axis represents the log_2_ fold change in gene transcription between the *B. abortus* WT and Δ*baaR* strains. Negative numbers represent genes downregulated and positive numbers genes upregulated in the Δ*baaR* background. The y-axis represents false discovery rate (FDR)-adjusted p-values transformed on a ‐log_10_ scale; a difference in gene transcription levels represented by ‐log_10_ FDR-adjusted p-values > 1 was considered significant. Genes with high fold changes and high FDR-adjusted p-values that were located in the same genomic regions were grouped. Genes present in the *bab2_0213-bab2_0217* locus are shown in red (*bab2_0213*: MFS transporter; *bab2_0214*: acyl-CoA dehydrogenase; *bab2_0216* (*acaD2*): pseudogene/enoyl-CoA hydratase-isomerase, *bab2_0217*: pseudogene/CoA-transferase family III); genes present in the *bab2_1036-bab2_1046* locus are shown in blue (*bab2_1036*: CoA-transferase family III, *bab2_1037-bab2_1040*: ABC transporter operon, *bab2_1041*: IclR, bab2_1042: pyruvate dehydrogenase E2 component, *bab2_1043*: periplasmic binding protein, *bab2_1044*: pseudogene/dehydrogenase E1 component/transketolase, *bab2_1045*: 3-hydroxyacyl-CoA dehydrogenase, *bab2_1046*: enoyl-CoA hydratase, *bab2_1047*: acyl-CoA dehydrogenase); and genes present in the *bab2_0277-bab2_0282* locus are shown in green (*bab2_0277*: glucose/methanol/choline oxidoreductase, *bab2_0278* and *bab2_0279*: branched-chain amino acid transport system permease proteins/ABC transporter, *bab2_0280* and *bab2_0281*: branched-chain amino acid transport system ATP-binding proteins/ABC transporter, *bab2_0282*: branched-chain amino acid transport system substrate-binding protein/ABC transporter). Other genes, highlighted in gray, were also strongly regulated, but no clear connection between those genes and the BaaR protein could be made (*bab1_0303*: UreG1 urease accessory protein*, bab1_0578:* BetI TetR transcriptional regulator, *bab1_0914:* DUF1127, *bab2_0548*: ABC transmembrane transporter). B) Genomic organization of the *bab2_0213-bab2_0217* operon. Genes *bab2_0213*/*bab2_0214*/*baaR* and genes *bab2_0216*/*bab2_0217* are on opposite strands and divergently transcribed. Gene *baaR* (black arrow) represses its own expression and that of the *bab2_0213-bab2_0217* genes.

*bab2_0217* encodes a protein with homology to CoA-transferase family III, a class of enzymes that catalyzes reversible transfer reactions of coenzyme A groups from CoA-thioesters to free acids (52). An early stop codon present in the sequence of this gene truncates the corresponding protein by 45 residues relative to related CoA-transferase family III proteins. The adjacent gene *bab2_0218* includes these 135 nucleotides missing from *bab2_0217;* the functional significance of *bab2_0217* truncation is not known.

Two additional gene sets were differentially regulated in *ΔbaaR* relative to WT. The first corresponds to 10 adjacent genes (*bab2_1036-bab2_1046)* (Figure 3A) with predicted enzyme functions similar to those associated with the *bab2_0213-bab2_0217* operon. Two of the most strongly upregulated genes, *bab2_1046* and *bab2_1045*, are annotated as enoyl-CoA hydratase/isomerase and 3-hydroxyacyl-CoA dehydrogenase, respectively, and may have functionally redundant roles with *bab2_0216*.Gene *bab2_1036* is annotated as a CoA-transferase family III protein and may be redundant with *bab2_0217*. Expression of the second set of genes, corresponding to *bab2_0277-bab2_0282* (Figure 3A) was significantly lower in *ΔbaaR* relative to WT. The majority of these genes encode components of an ATP-binding-cassette (ABC) transport system predicted to be involved in branched chain amino acid transport. Comparison of the amino acid sequence of the cognate periplasmic binding protein (PBP) with the sequences of PBPs that co-crystallize with *bona fide* ligands revealed that Bab2_0282 shares 54% identity with a *Burkholderia mallei* PBP (Protein Data Bank [PDB]: 3I09) that co-crystallizes with acetoacetate. Residues involved in the interaction with acetoacetate in this PBP are conserved in Bab2_0282 (Figure S2), suggesting that the *B. abortus bab2_0278-bab2_0282* ABC transporter operon is involved in uptake of acetoacetate or a related molecule. Acetoacetate is produced in the liver during ketogenesis. Under certain conditions, the acetyl-CoA formed in the liver from β-oxidation of fatty acids can be converted into ketone bodies (acetoacetate, D-β-hydroxybutyrate, and acetone) for export to other tissues (53,54). This *Brucella* ABC transporter operon is proximal to a gene (*bab2_0277*)annotated as a glucose/methanol/choline oxidoreductase, which is also expressed at significantly lower levels in *ΔbaaR*. In *B. mallei*, a similar gene (*BMA2933*) also co-occurs with the related ABC transporter, suggesting that this system may have a function in ketone metabolism.

Finally, genes *bab1_0303* (UreG1 urease accessory protein), *bab1_0578* (BetI TetR transcriptional regulator), *bab1_0914* (DUF1127), and *bab2_0548* (ABC transmembrane transporter) also showed significant differential regulation in our data set. The functional significance of these transcriptional changes is not known (see Table S2 for the full data set).

### BaaR binds palindromic motifs in its own promoter region

To evaluate the mechanism of BaaR-dependent transcriptional repression, we measured binding between purified BaaR and a region of DNA corresponding to its promoter. Since the IclR protein family recognizes and specifically interacts with palindromic motifs (11), the DNA sequence corresponding to the promoter region between *baaR* and *bab2_0216* was analyzed for palindromes. We identified two similar palindromic regions that were 44 and 35 nucleotides away from the *bab2_0216* and *baaR* start codons, respectively. These two regions were named BaaR binding site 1 (BBS1) and BaaR binding site 2 (BBS2), and each consisted of two overlapping palindromic sequences (Figures 4 and 5). A 375-nucleotide fluorescent DNA probe corresponding to the promoter region between *baaR* and *bab2_0216* was used for gel shift experiments. As expected, 5 ng of this “long” fluorescent DNA probe corresponding to the WT sequence (L^WF^: long, wild-type, fluorescent) was shifted in size on the gel in the presence of increasing concentrations (2-500 nM) of purified BaaR, confirming that BaaR interacts with the DNA region between *bab2_0215* and *bab2_0216* (Figure 4A, (1)).

**Figure 4:**
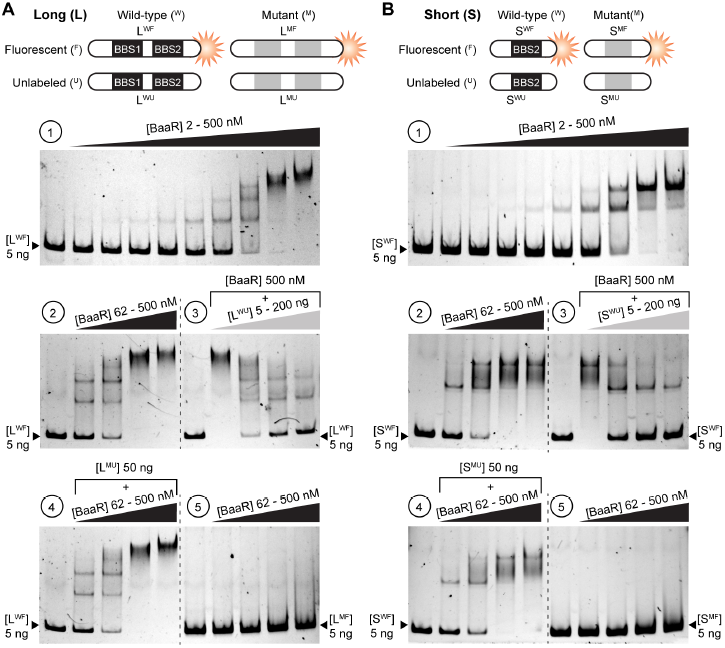
Electrophoretic mobility shift assay of the interaction between purified BaaR protein (residues 9-283) and the *bab2_0215-bab2_0216* intergenic region. A) (1) A long WT fluorescent DNA probe (L^WF^, 375 nucleotides long) corresponding to the intergenic region between *baaR* and *bab2_0216*, including BBS1 and BBS2, was mixed with increasing concentrations (2, 4, 8, 16, 31, 62, 125, 250, and 500 nM) of purified BaaR and run on a 5% native acrylamide gel. Lane 1 represents 5 ng of L^WF^ alone (i.e., without BaaR) and was used as a size control. (2) The L^WF^ (5 ng) was mixed with increasing concentrations (62, 125, 250, and 500 nM) of purified BaaR and run on a 5% native acrylamide gel. Control lane 1 represents 5 ng L^WF^ alone (i.e., without BaaR). (3) The L^WF^ (5 ng) was mixed with 500 nM purified BaaR and increasing concentrations (5, 50, 100, and 200 ng) of a long WT unlabeled DNA probe (L^WU^), corresponding to the intergenic region between *baaR* and *bab2_0216*, including BBS1 and BBS2. Control lane 1 represents 5 ng L^WF^ alone (i.e., without BaaR). (4) The conditions used in this assay were similar to those used in (2), except that 50 ng of the L^MU^ DNA probe (long mutated and unlabeled), corresponding to the intergenic region between *baaR* and *bab2_0216* and carrying mutated BBS sequences, were added. Control lane 1 corresponds to 5 ng L^WF^ alone (i.e., without BaaR). (5) The conditions used in this assay were similar to those used in (2), except that 5 ng L^MF^ probe (long mutated and fluorescent) was used instead. Control lane 1 corresponds to 5 ng L^MF^ alone (i.e., without BaaR) and was used as a size reference. B) The gel shift assays in (1), (2), (3), (4), and (5) were performed as described in (A); however, instead of using a DNA probe targeting the full-length *baaR*-*bab2_0216* intergenic region, a shorter (196 nucleotides long) WT fluorescent DNA probe (S^WF^) targeting only half of this region, carrying BBS2 but not BBS1, was used. A cartoon representation of the different DNA probes used in this assay is presented at the top of each panel.

To determine if this interaction was specific, we conducted a series of control experiments. As previously observed, 5 ng of the L^WF^ probe were shifted in size when mixed with increasing concentrations (62-500 nM) of BaaR (Figure 4A, (2)). However, this shift was weakened when 5 ng of the L^WF^ probe were mixed with 500 nM BaaR, a concentration at which all of the DNA probe was shifted, and increasing concentrations (5-200 ng) of an equivalent unlabeled (non-fluorescent) DNA probe (L^WU^: long, wild-type, unlabeled), indicating that the fluorescent and unlabeled probes competitively interacted with BaaR (Figure 4A, (3)). This result suggests that the interaction between BaaR and its promoter is specific. In a related control, 5 ng of the L^WF^ DNA probe and 50 ng of an BBS-mutated unlabeled DNA probe (L^MU^: long, mutated, unlabeled) were mixed together. This mutated probe did not disrupt the BaaR/L^WF^ interaction, suggesting BaaR binding requires the BBS1 and BBS2 regions (Figure 4A, (4)). Finally, when we performed a gel shift assay with a fluorescent DNA probe carrying mutated BBS1 and BBS2 (L^MF^: long, mutated, fluorescent), no shift was observed, even at the highest BaaR concentrations (Figure 4A, (5)). We conclude that BaaR constitutively and specifically interacts with the *bab2_0215-bab2_0216* promoter region, and this interaction requires BBS1 and BBS2. Equivalent gel shift assays were conducted using a shorter DNA probe (196 nucleotides) harboring only one BBS (BBS2) (Figure 4B). This probe behaved exactly the same as the long probe in the presence of the purified BaaR (Figure 4B). Control experiments performed with the different unlabeled or mutated DNA probes also confirmed the specificity of this interaction (Figure 4B). As discussed above, BaaR contains approximately 38 additional amino acids at its N-terminus. To assess the role of this N-terminal extension in the interaction between BaaR and its target DNA, we performed gel shift assays using a BaaR mutant protein missing these additional N-terminal amino acids. We observed no significant differences in probe shift between the short (residues V40-P284) and WT BaaR proteins; both resulted in shifts of the long (289 nucleotides, contains BBS1 and BBS2) and short (196 nucleotides, contains BBS2) fluorescent DNA probes (Figure S3).

### The palindromic regions between *baaR* and *bab2_0216* are required for recognition by BaaR

To identify the motifs in the BBS region between *baaR* and *bab2_0216* required for interaction with BaaR, each palindrome of BBS2 was independently mutated. We then measured binding of the corresponding 196-nucleotide fluorescent DNA probe to BaaR by gel shift assay. In comparison to the WT BBS2 containing probe, no gel shift was observed when the first half of the first palindromic motif (region 1) was mutated, suggesting no interaction between the DNA probe and BaaR (Figures 5A and 5B). Mutation of BBS2 region 2, corresponding to overlapping region between the first and second palindromic motifs, also ablated binding of the probe to BaaR (Figures 5A and 5B). Mutation of the second half of the second palindrome (region 3) resulted in an intermediate effect; a partial gel shift of the corresponding DNA probe was still observed with the highest BaaR concentrations (250 and 500 nM) (Figure 5A and 5B). As expected, mutating regions 1, 2 and 3 completely ablated a gel shift (Figure 5B). We conclude that regions 1 and 2, present in the TTTCGC/GCGAAA palindrome, are required for interaction with BaaR, and that region 3 plays a minor role in the BaaR/DNA interaction. We note that DNA regions upstream of *bab2_0277-bab2_0282* and *bab2_1036-bab2_1046* do not contain similar palindromic sequences recognized by BaaR. This suggests that the transcriptional regulation of these loci observed in our RNA-seq data is indirect.

**Figure 5:**
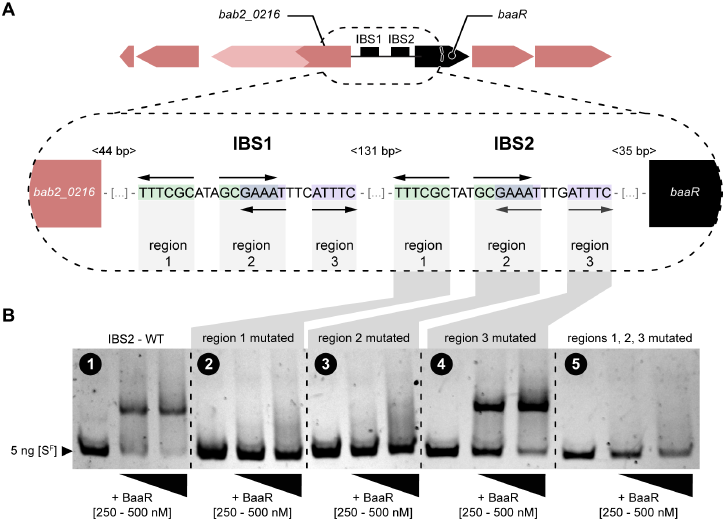
Electrophoretic mobility shift assay of the interaction between purified BaaR protein and different DNA probes mutated in the BBS2 region. A) Cartoon representation of the intergenic region between *baaR* and *bab2_0216*. The sequences of the two similar BaaR binding sites (BBS1 and BBS2), consisting of two overlapping palindromes, are presented. The first palindrome is the TTTCGC/GCGAAA sequence and the second palindrome is the GAAAT/ATTTC sequence. Each BBS can be divided into three regions, corresponding to the different palindromic sequences. The BBS1-BBS2, BBS2-*baaR* and BBS1-*bab2_0216* nucleotide distances are shown in light gray. B) Gel shift assay of the interaction between the BaaR protein (250 and 500 nM) and 5 ng of a short fluorescent DNA probe (S^F^, 196-nucleotides long) carrying different mutations of BBS2 regions. ESMA performed with (1) wild-type BBS2, (2) BBS2 palindromic region 1 mutated, (3) BBS2 palindromic region 2 mutated, (4) BBS2 palindromic region 3 mutated, and (5) BBS2 palindromic regions 1, 2 and 3 mutated. In each panel, lane 1 (the control) corresponds to 5 ng of the different S^F^ probes alone (i.e., in the absence of the BaaR protein).

### BaaR interacts with BBS1 and BBS2 *in vivo* and regulates transcription of the *bab2_0213-bab2_0217* operon

We next evaluated whether BaaR interacts with the *baaR-bab2_0216* promoter region *in vivo*, and whether both BBSs are required for transcriptional regulation in *B. abortus*. We transformed WT and Δ*baaR* strains with a plasmid carrying a transcriptional fusion of the intergenic region between *bab2_0215* and *bab2_0216* to *lacZ*. This *lacZ* transcriptional reporter construct (P*_baaR_-lacZ*) contains the *baaR* promoter region including both BBS1 and BBS2.

In the WT strain, β-galactosidase activity was low, suggesting constitutive repression of transcription by BaaR. In the *ΔbaaR* strain, β-galactosidase activity was increased by a factor of 10 relative to WT, suggesting that deletion of *baaR* derepresses transcription under the control of its own promoter (Figure 6A, (1)). Surprisingly, mutation of BBS1 and BBS2 (BBS1^mut^/BBS2^mut^) resulted in low β-galactosidase activity in both strains. It is likely that these tandem BBS1 and BBS2 mutations disrupt the ability of RNA polymerase to induce transcription from this promoter (Figure 6A, (1)). Mutation of BBS1 alone (BBS1^mut^/BBS2^WT^) had little effect on BaaR-dependent transcriptional repression: β-galactosidase activity was low in the WT and high in the *ΔbaaR* strain (Figure 6A, (1)). However, in the WT strain, the measured β-galactosidase activity was higher from the BBS1^mut^/BBS2^WT^ than from the BBS1^WT^/BBS2^WT^ reporter. This indicates that the integrity of both BBSs is likely required for efficient repression by BaaR, perhaps by permitting multiple BaaR dimers to interact simultaneously. Mutation of BBS2 alone (BBS1^WT^/BBS2^mut^) resulted in low β-galactosidase activity in both the WT and *ΔbaaR* strains, further supporting that RNA polymerase cannot efficiently induce transcription from this sequence (Figure 6A, (1)). We conclude that the integrity of BBS2 is essential for the proper interaction of transcription factors at this promoter.

**Figure 6:**
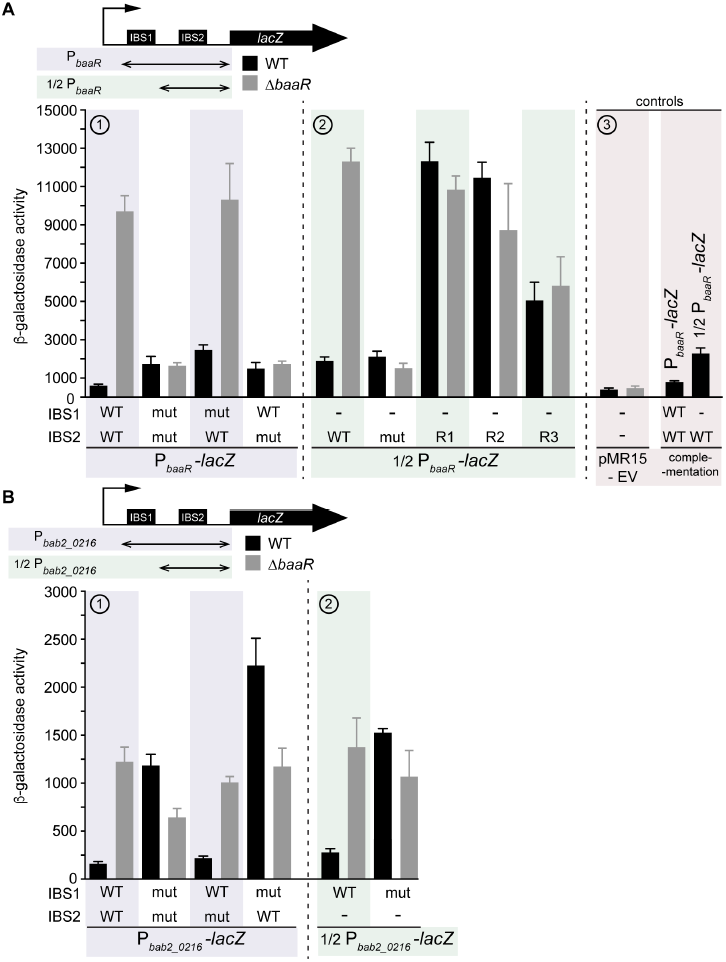
β-galactosidase activities of the *B. abortus* WT and *B. abortus* Δ*baaR lacZ* reporter strains. Error bars represent standard deviations A) (1) β-galactosidase activities of WT and Δ*baaR B. abortus* strains transformed with different versions of the P*_baaR_-lacZ* reporter gene (boxed in violet). These reporter genes are a transcriptional fusion between *lacZ* and the *baaR-bab2_0216* intergenic region encompassing WT or mutant (mut) BBS1 and BBS2. (2) The WT and Δ*baaR B. abortus* strains were transformed with different versions of the 1/2P*_baaR_-lacZ* reporter gene (boxed in green). These reporter genes corresponded to a transcriptional fusion between *lacZ* and half of the *baaR-bab2_0216* intergenic region carrying the WT or mutated BBS2. BBS2 regions 1 (R1), 2 (R2), and 3 (R3) were also mutated individually. (3) Negative controls (boxed in pink): the WT and Δ*baaR B. abortus* strains were transformed with the pMR15 empty vector (EV) and a Δ*baaR* strain was complemented with the WT *baaR* allele and transformed with P*_baaR_-lacZ* and 1/2P*_baaR_-lacZ* reporter genes. B) As in (A), the WT and Δ*baaR B. abortus* strains were transformed with different versions of the (1) P*_bab2_0216_*-*lacZ* (boxed in violet) or (2) 1/2P*_bab2_0216_*-*lacZ* (boxed in green) reporter genes. These reporter genes correspond to a transcriptional fusion between *lacZ* and the reverse complement of either the full-length (encompassing BBS1 and BBS2) or half (BBS1 only) of the *baaR-bab2_0216* intergenic region.

When half of the intergenic region present between *baaR* and *bab2_0216* was fused to *lacZ*, the corresponding reporter gene (1/2P*_baaR_*-*lacZ*) behaved similarly to the BBS1^mut^/BBS2^WT^ reporter: β-galactosidase activity was low in the WT and high in the *ΔbaaR* strain (Figure 6A, (2)). Again, transcription from the corresponding reporter with only one BBS (BBS2^WT^) was not as repressed as that from the reporter containing both BBS sequences (BBS1^WT^/BBS2^WT^) (Figure 6A, (2)). Mutation of BBS2 (BBS2^mut^) resulted in low β-galactosidase activity in both strains and could be attributed to a possible disruption in transcription (Figure 6A, (2)). To overcome this problem, we mutated smaller regions of BBS2. When BBS2 region 1 (BBS2^R1^) or 2 (BBS2^R2^) was mutated, the corresponding β-galactosidase activity was high in both the WT and mutant strains (Figure 6A, (2)). From these data, we conclude that the integrity of the transcription initiation site is preserved in these constructs. This result also confirmed that BaaR in the WT strain fails to interact with BBS2^R1^ or BBS2^R2^. When region 3 (BBS2^R3^) was mutated, intermediate β-galactosidase activity was measured in both strains (Figure 6A, (2)), which confirmed our previous in vitro observations (Figure 5B). Taken together, these results suggest that BBS2 is required for BaaR-dependent transcriptional regulation of *bab2_0215*, *bab2_0214*, and *bab2_0213*. However, the presence of BBS1 ensures even greater transcriptional repression. WT or Δ*baaR* strains carrying the empty vector (pMR15) or the *baaR* complemented strain were used as controls (Figure 6A, (3)).

We next evaluated whether transcription from *bab2_0216-bab2_0217* exhibited a similar transcriptional profile to *baaR*. We constructed a new *lacZ* transcriptional reporter in which the reversed and complemented intergenic sequence between *baaR* and *bab2_0216* was fused to *lacZ* (P*_bab2_0216_*-*lacZ*). This construct was transformed into the WT or Δ*baaR* strains. β-galactosidase activity in Δ*baaR* was fivefold higher than that in WT, providing evidence that BaaR represses transcription from this reporter as well (Figure 6B, (1)). The β-galactosidase activity observed under this reporter was generally lower than that under P*_baaR_-lacZ*, suggesting weaker transcription from this promoter. When BBS1 and BBS2 were mutated simultaneously (BBS1^mut^/BBS2^mut^), both the WT and Δ*baaR* strains exhibited increased β-galactosidase activity (Figure 6B, (1)). However, in the WT strain, these activities were higher and comparable with the β-galactosidase activity levels measured in the Δ*baaR* strain carrying the BBS1^WT^/BBS2^WT^ reporter gene. In both strains, mutation of BBS2 only (BBS1^WT^/BBS2^mut^) had no effect on β-galactosidase activity, whereas mutation of BBS1 only (BBS1^mut^/BBS2^WT^) induced greater activity (Figure 6B, (1)).

Lastly, a reporter containing half of the *bab2_0216* promoter region was evaluated (1/2P_bab2_0216_-*lacZ*) (Figure 6B, (2)). Transcriptional activity under this reporter in the WT strain was fivefold lower than that in the Δ*baaR* strain and was comparable with that in the BBS1^WT^/BBS2^mut^ reporter strains (Figure 6B, (2)). When BBS1 was mutated, activity increased in both strains and was comparable with that in the BBS1^mut^/BBS2^mut^ reporter strains (Figure 6B, (2)).

Together, these results provide evidence that BBS1 is required for BaaR-dependent transcriptional regulation of *bab2_0216* and *bab2_0217*. BBS1 and BBS2 mutations did not disrupt the ability of RNA-polymerase to induce transcription from this promoter, suggesting that the *bab2_0216-bab2_0217* initiation site does not overlap with BBS1 or BBS2.

### ε-aminocaproic acid derepresses transcription from a BaaR-regulated promoter

BaaR, like other IclR proteins, has a C-terminal LBD with a cavity accommodating small molecules. Interaction with specific molecules can positively or negatively modify the affinity of the protein for its target DNA (11,14,17,41,42,44,55). We screened for small molecules that affect transcription from a BaaR-regulated reporter plasmid. Specifically, we transformed WT *Brucella ovis*, a closely related Biosafety level 2 (BSL2) surrogate for *B. abortus,* with the P*_baaR_-lacZ* reporter and inoculated this strain into 96-well plates containing 480 distinct, individual small molecules. A single molecule, ε-aminocaproic acid, activated transcription from the P*_baaR_-lacZ* reporter under this cultivation condition (Figure 7A). We confirmed this hit in a *B. abortus* strain carrying the same reporter plasmid (Figure 7B). ε-aminocaproic acid is a six-carbon molecule with a carboxyl and amine group. It is a lysine derivative and analog used in clinical settings to promote blood clotting (56,57). In this same transcription induction screen, acetate and acetoacetate did not affect transcription (Table S3). We thus conclude that neither acetoacetate nor acetate are ligands that induce transcription, even though we observe acetate bound to the LBD in the BaaR crystal structure.

**Figure 7:**
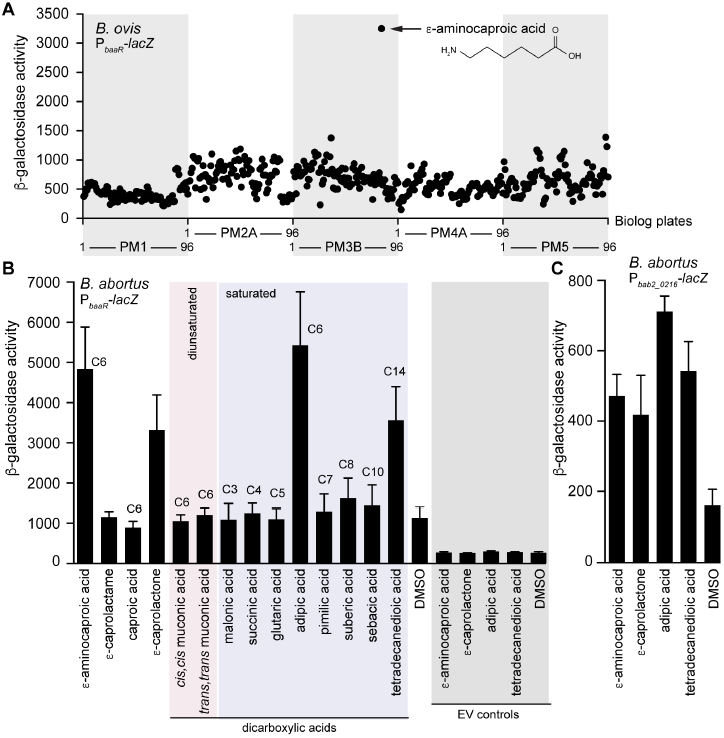
Screening of small molecules to identify an BaaR effector. A) A WT *B. ovis* strain carrying the P*_baaR_-lacZ* reporter gene was grown overnight (37°C, 5% CO_2_) on Phenotype MicroArrays (Biolog), and β-galactosidase activity was measured under each condition. For each plate (PM1, PM2A, PM3B, PM4A, and PM5), molecules 1-96 are delimited on the x-axes. Molecule G9 on plate PM3B, corresponding to ε-aminocaproic acid, is indicated with a black arrow, and its molecule structure is shown. A list of the different growth conditions evaluated and the corresponding β-galactosidase activities measured can be found in Table S3. B) A WT *B. abortus* strain carrying the P*_baaR_-lacZ* reporter was grown overnight in the presence of 4 mM of the indicated molecules or DMSO. Diunsaturated and saturated dicarboxylic acids are highlighted by a pink or violet rectangle, respectively. As negative controls (gray rectangle), *B. abortus* carrying the pMR15 empty vector control (EV) was grown in the presence of 4 mM ε-aminocaproic acid, adipic acid, tetradecanedioic acid, ε-caprolactone or DMSO. Error bars represent standard deviations C) As in (B), a WT *B. abortus* strain carrying the P*_bab2_0216_*-*lacZ* reporter gene was grown overnight in the presence of 4 mM ε-aminocaproic acid, ε-caprolactone, adipic acid, tetradecanedioic acid, or DMSO. Error bars represent standard deviations.

### Adipic acid, tetradecanedioic acid, and ε-caprolactone also derepress transcription

Given the results of our initial screen, we evaluated other related small molecules for their ability to activate transcription from a BaaR-regulated reporter. Selection of these molecules was based on their metabolic and physiological properties and chemical and structural similarities shared with ε-aminocaproic acid (Figure S4). To narrow the number of molecules to be tested, the sequence of BaaR was compared with those of IclR proteins previously described to regulate similar metabolic pathways. BaaR shares 59% and 54% identity with DcaS (ACIAD1684) and DcaR (ACIAD1688), two *Acinetobacter* IclR proteins potentially involved in the transcriptional regulation of a *dca* β-oxidation operon (36,37,58). BaaR also has high identity (66%) to the *Pseudomonas* DcaR protein (PA1630) involved in regulation of ε-caprolactam catabolism and β-oxidation (36,38). In *Acinetobacter* ADP1 and *P. aeruginosa*, *dca* operons have been previously described as essential for growth on adipic acid or ε-caprolactam as the sole carbon sources (36,38). Interestingly, a *cis,cis* muconic transporter (ACIAD1681) is proximal to the *Acinetobacter dca* operon (39); this transporter is 67% identical to Bab2_0213 in *B. abortus*. The ability of adipic acid, ε-caprolactam, and *cis,cis*, muconic acid to derepress transcription of P*_baaR_-lacZ* was therefore investigated in *B. abortus*. Addition of *cis,cis* muconic acid did not enhance transcription, but 4 mM of adipic acid strongly activated transcription (Figure 7B and Figure S4). To evaluate the specificity of this activation, shorter or longer dicarboxylic acids (C3 to C14) were assessed as well. Only tetradecanedioic acid, a C14 dicarboxylic acid, significantly derepressed transcription from our reporter (Figure 7B and S4).

*Trans,trans* muconic acid, a dicarboxylic acid closely related to *cis,cis* muconic acid, was also evaluated, but it had no effect on transcription from the reporter (Figures 7B and S4). Interestingly, the six-carbon fatty acid caproic acid had no effect on BaaR-dependent repression of P*_baaR_-lacZ* reporter gene transcription, while its cyclic form, ε-caprolactone (59), significantly derepressed transcription (Figure 7B and S4). Conversely, ε-caprolactam, a cyclic ε-aminocaproic acid (60), had no effect on transcription (Figure 7B and S4). As a control, we cultivated the reporter strain in the presence of DMSO, which was used to solubilize most of the small molecules evaluated. Transcription from this reporter was unchanged by addition of the DMSO (Figure 7B). We also confirmed that the different molecules had no effect on an empty vector control strain (Figure 7B). All molecules were also assessed in a *B. abortus* strain carrying the P_*bab2_02i6*_*-lcicZ* reporter. β-galactosidase activity was significantly enhanced in the presence of 4 mM ε-aminocaproic acid, adipic acid, tetradecanedioic acid, and ε-caprolactone (Figure 7C).

### Evidence that adipic acid is transported by Bab2_0213, an MFS transporter

We next evaluated whether the molecules found to induce transcription from a BaaR-dependent reporter *(P_*baaR*_*-lacZ*)* are specifically transported by the MucK-like transporter Bab2_0213. We also tested whether Bab2_0214 (acyl-CoA dehydrogenase), Bab2_0216 (enoyl-CoA hydratase/isomerase and 3-hydroxyacyl-CoA dehydrogenase), and Bab2_0217 (CoA-transferase family III) affect BaaR-regulated transcription. We generated in-frame deletions of each of these genes in *B. abortus* and transformed the corresponding null mutant strains with the P_*baaR*_-*lacZ* reporter plasmid. Each deletion strain was grown in the presence of increasing concentrations of the different activating molecules, and β-galactosidase activity was measured under each condition.

Transcription reporter activity in all four null mutant strains was the same as that in the WT in the presence of ε-aminocaproic acid, tetradecanedioic acid, or ε-caprolactone (Figure 8A-C). However, the strain harboring the *bab2_0213* deletion exhibited significantly lower transcriptional activity than the WT strain and other deletion strains in the presence of adipic acid (Figure 8D). Complementation of *Δbab2_0213,* by re-introducing a WT copy of *bab2_0213,* restored the WT transcriptional phenotype (Figure 8E). We conclude that, at the concentrations tested, Bab2_0213 is involved in transport of adipic acid. It is not known how ε-aminocaproic acid, tetradecanedioic acid, or ε-caprolactone is transported. We further conclude that deletion of *bab2_0214*, *bab2_0216*, or *bab2_0217* did not affect the response of *B. abortus* to these inducing molecules. This provides evidence that none of these enzymes are required for BaaR-dependent transcriptional regulation by adipic acid, ε-aminocaproic acid, tetradecanedioic acid, or ε-caprolactone.

**Figure 8:**
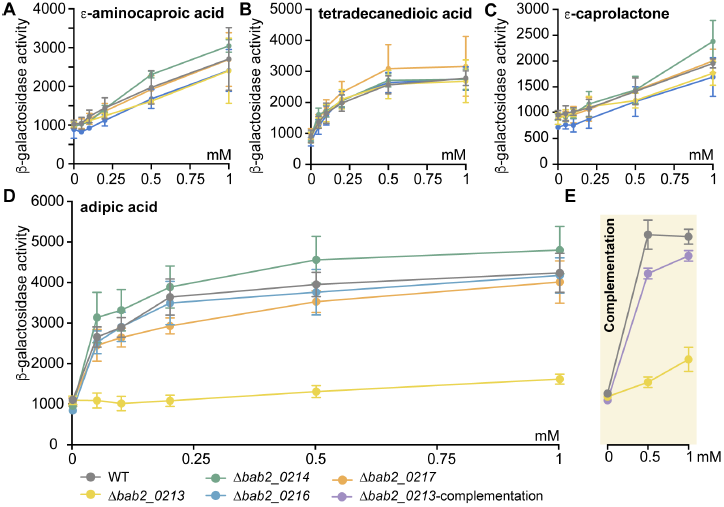
Effects of ε-aminocaproic acid, ε-caprolactone, adipic acid, and tetradecanedioic acid on transcriptional activation of a BaaR-repressed reporter gene in different *B. abortus* deletion backgrounds. The P*_baaR_-lacZ* reporter gene was transformed into the WT (gray line), Δ*bab2_0213* (yellow line), Δ*bab2_0214* (green line), Δ*bab2_0216* (blue line), and Δ*bab2_0217* (orange line) strains. Each strain was grown overnight in the presence of 0, 0.05, 0.1, 0.2, 0.5, or 1 mM of A) ε-aminocaproic acid, B) tetradecanedioic acid, C) ε-caprolactone, or D) adipic acid. Right panel (light yellow): the Δ*bab2_0213* strain was complemented (purple line) with a WT copy of *bab2_0213*, and its ability to transport and respond to 0, 0.5, and 1 mM adipic acid was evaluated and compared with those of the WT (gray line) and Δ*bab2_0213* (yellow line) strains. Error bars represent standard deviations.

### Adipic acid, ε-aminocaproic acid, tetradecanedioic acid, and ε-caprolactone do not interact directly with BaaR

Given the ability of particular organic acids to derepress transcription from a BaaR-dependent reporter *in vivo*, we next evaluated whether adipic acid, ε-aminocaproic acid, tetradecanedioic acid, or ε-caprolactone modify BaaR binding to its target DNA. We mixed a fluorescent DNA probe corresponding to the BBS2 region with purified BaaR and (separately) 1 mM of each small molecule. We conducted this experiment using a range of BaaR concentrations.

None of the molecules tested had any effect on BaaR binding to the probe (Figure 9A-E). However, the possibility that these molecules interact with the BaaR LBD without affecting the interaction of BaaR with DNA could not be ruled out in this assay. Therefore, we also performed isothermal titration calorimetry (ITC) measurements, which allow characterization of the affinities between IclR proteins and small molecules at micromolar levels (41,55,61). Specifically, we performed ITC measurements between purified BaaR LBD and adipic acid, which is likely transported by Bab2_0213 and thus a *bona fide* direct activating signal. We observed no interaction between adipic acid and the LBD of BaaR, suggesting an indirect effect of adipic acid *in vivo* (Figure 9F).

**Figure 9:**
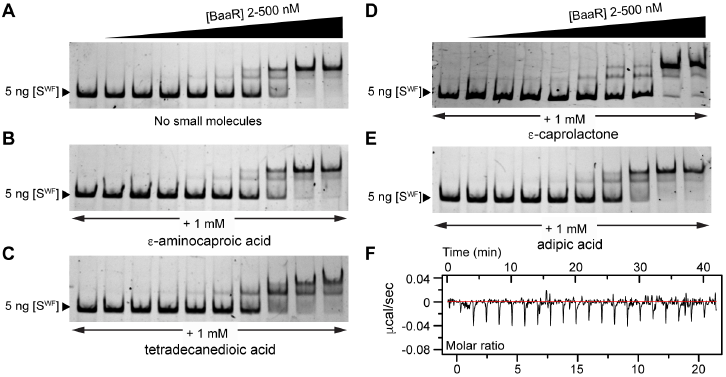
Electrophoretic mobility shift assays were performed with purified BaaR protein and a short WT fluorescent DNA probe (S^WF^, 196-nucleotide long) carrying BBS2 in the A) absence or B) presence of 1 mM ε-aminocaproic acid, C) tetradecanedioic acid, D) ε-caprolactone, or E) adipic acid. The S^WF^ DNA probe (5 ng) was mixed with increasing concentrations (2, 4, 8, 16, 31, 62, 125, 250, and 500 nM) of BaaR protein and 1 mM of small molecule. Samples were run on a 5% native acrylamide gel. F) Isothermal titration calorimetry performed using the purified BaaR LBD and adipic acid; 1 μl 10 mM adipic acid solution was injected into a cell containing 200 μl of a 50 μM protein solution. A total of 20 injections were performed.

### Deleting *baaR* does not affect *B. abortus* intracellular entry or replication

As outlined earlier, the general stress response sigma factor σ^E1^ indirectly activates transcription of BaaR. Given the known role of σ^E1^ in *B. abortus* infection biology (4,7), we evaluated whether BaaR contributes to *B. abortus* infection *in vitro*. Cultured THP-1 macrophage-like cells were infected (multiplicity of infection = 100) with WT and *ΔbaaR B. abortus* strains. No significant differences in WT and *ΔbaaR* CFU recovered from infected THP-1 cells were observed at any time point evaluated (1, 24, and 48 hours post infection), suggesting that BaaR and *bab2_0213-bab2_0217* repression are not required for entry or replication in a THP-1 *in vitro* infection model (Figure 10). This result is consistent with a previous study in which *Brucella melitensis* 16M with deletion of a gene orthologous to *baaR, BMEII1022,* was not attenuated in mouse spleen colonization at 1 week post infection (62).

**Figure 10:**
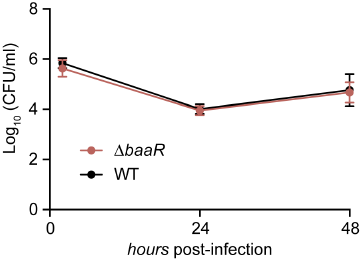
*In vitro* infection assay. Infection of inactivated THP-1 cells with WT *B. abortus* strain 2308 (black line) and Δ*baaR* (red line). The number of *B. abortus* CFUs recovered from the THP-1 cells at 1, 24, and 48 hours post infection is plotted. Error bars represent standard deviations.

## Discussion

As a facultative intracellular pathogen, *B. abortus* is exposed to many different microenvironments during its lifecycle. The general stress response sigma factor, σ^E1^, controls the transcription of dozens of genes and is required for *B. abortus* survival under stress and chronic infection conditions in mouse disease models (4,5,7,9,10). In this manuscript, we present structural and functional studies of a transcriptional regulator of the IclR family, BaaR, which is indirectly activated by σ^E1^. BaaR strongly represses transcription from two divergently transcribed operons (Figures 3 and 6) homologous to the *dca* β-oxidation operons of *Acinetobacter* spp. and *Pseudomonas* spp., which have been implicated in growth on adipic acid or **e**-caprolactam as a sole carbon source (36-;38) (Figure 11). In *Acinetobacter* and *Pseudomonas,* it has been postulated that IclR transcriptional regulators control *dca* operon expression, although this hypothesis has not been tested experimentally. When compared to other IclR proteins, BaaR is closely related to *Acinetobacter* DcaR (ACIAD1688; 54% identity) and DcaS (ACIAD1684; 59% identity) and *Pseudomonas* DcaR (PA1630; 66% identity) (Figure 11). All four of these proteins possess an N-terminal extension in comparison to other IclR proteins, and very similar LBD cavities (Figures 2C and S1), suggesting they recognize structurally related small molecules. The function of the extended N-termini in these related proteins is unclear, and deletion of the N-terminus of BaaR did not affect DNA binding *in vitro* (Figure S3). It is possible that in a cellular context, this structural region affects protein-DNA interactions or protein stability in the presence of specific molecular signals. *In vivo*, BaaR represses transcription of the *bab2_0213-bab2_0217* locus (including its own gene: *bab2_0215*). Adipic acid and other related organics relieve this repression, resulting in initiation of *bab2_0213-bab2_0217* transcription. It remains unclear how transcription of this operon is maintained despite increasing concentrations of BaaR in the cell. It is conceivable that like the TtgV protein of *P. putida* (63,64) and other proteins belonging to the IclR family (14,15,65,66), reduced affinity of BaaR for DNA upon ligand binding is sufficient to overcome the effects of increased concentrations in the cell.

**Figure 11:**
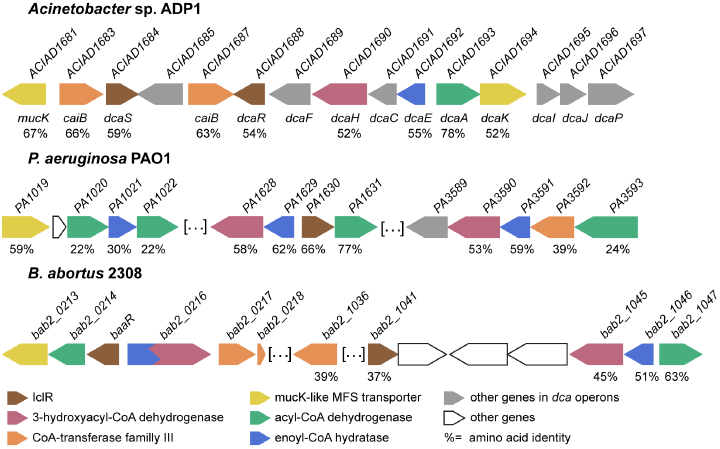
Genomic context and protein sequence identity of the β-oxidation operons in *Acinetobacter*, *Pseudomonas*, and *B. abortus*. A) Genes annotated as IclR proteins are in brown, 3-hydroxyacyl-CoA dehydrogenases in red, CoA-transferases family III in orange, MucK-like transporters in yellow, acyl-CoA dehydrogenases in green, and enoyl-CoA isomerases/hydratases in blue. Other genes present in the *Acinetobacter dca* operon that are not regulated by BaaR in *B. abortus* are in gray. Genes not related to β-oxidation and present in a different locus are shown in white. Gene annotation numbers are presented above each gene. Below each gene, the percentage of identity between the corresponding protein and the equivalent Bab2_0213-Bab2_0217 protein is presented.

When compared to the *Acinetobacter dca* locus, the *B. abortus bab2_0213-bab2_0217 dca*-like operon (regulated by BaaR) appears incomplete. Indeed, genes essential for the β-oxidation of dicarboxylic acids in *Acinetobacter* are degenerate or absent in *B. abortus* (Figure 11). It is therefore difficult to conclude that *bab2_0213-bab2_0217* is truly involved in β-oxidation metabolism. The RNA-seq studies presented here identified additional genes, present in the *bab2_1036-bab2_1046* locus, that are strongly upregulated after deletion of *baaR.* Based on sequence homology, these genes are paralogs and likely have functions that overlap or complement *bab2_0214-bab2_0217* (Figures 3 and 11). It is conceivable that like *Pseudomonas* PAO1, multiple clusters of *dca*-like genes have redundant functions in dissimilation of short‐ or medium-chain dicarboxylic acids (Figure 11) (36). No transporter related to *bab2_0213* was found proximal to this set of paralogous genes or in *B. abortus* genome, suggesting that *bab2_0213-bab2_0217* and parlalogs could process a shared substrate that is transported by Bab2_0213. Bab2_0213 has high sequence identity with MucK-like MFS transporters located proximal to the *Acinetobacter* (*ACIAD1681 (mucK)* and *ACIAD1694 (dcaK)*; 67% and 52%) and *Pseudomonas (PA1019;* 59%) *dca* operons. In *Acinetobacter,* MucK is essential for growth in presence of *cis, cis* muconic acid but not adipic acid. A potential role for DcaK in adipic acid transport has been discussed, although it has never been experimentally investigated (Figure 11) (36,38,39). Our results provide evidence that Bab2_0213 is involved in adipic acid transport, though we cannot rule out the possibility that this protein could also transport other types of molecules. Homology between Bab2_0213 and MFS transporters associated with the *dca* genes in *Acinetobacter* or in *Pseudomonas* clearly suggests that the molecules transported by these systems are structurally related to adipic acid or *cis,cis*-muconate.

The RNA-seq analysis of WT *B. abortus* and the *ΔbaaR* strain also revealed a predicted ABC transport system *(bab2_0277-bab2_0282)* that was indirectly activated by BaaR. The primary structure of the periplasmic binding protein (PBP) in this system (Bab2_0282) is 54% identical to a *Burkholderia mallei* PBP that co-crystallized with bound acetoacetate (PDB ID: 3I09). Residues involved in this interaction with acetoacetate are also present in Bab2_0282, suggesting that Bab2_0282 may transport acetoacetate or a closely related molecule. In mammals, acetyl-CoA formed in the liver during fatty acid β-oxidation can be converted into acetoacetate and released into the bloodstream as an energy source during periods of starvation or intense physical activity (53,54). We have no evidence that exogenous acetoacetate is actively transported by *B. abortus,* nor do we understand the metabolic connection (if any) to the *bab2_0213-bab2_0217* locus. However, it is conceivable that this system could be involved in utilizing acetoacetate or related substrates produced by the host.

In conclusion, we have demonstrated that adipic acid and the related molecules ε-aminocaproic acid, tetradecanedioic acid, and ε-caprolactone activate transcription of the *bab2_0213-bab2_0217* locus in *B. abortus,* though these molecules do not activate transcription through a direct interaction with BaaR. This suggests that the actual activating molecule *in vivo* is either a metabolic product of these compounds, or that these inducing compounds activate some undefined metabolic pathway involved in the synthesis of the BaaR-activating signal. The possible relevance of such molecules in the life cycle of *B. abortus* and the functional significance of *baaR* transcriptional activation by σ^E1^ remain undefined. Adipic acid is produced by oxidation of fatty acids but is not naturally abundant. We note that very little is known about the *B. abortus* life cycle outside the host. In the wild, *B. abortus* can persist for weeks in aborted fetuses, a major source of contagion (67). However, *B. abortus* can also persist for weeks in soil or on vegetation. How *B. abortus* survives in these harsh and competitive environments is unknown, and it is possible that the genes investigated in this study enable *B. abortus* to metabolize unusual substrates found outside the mammalian host. To conclude, we propose a model (Figure 12) whereby dimeric BaaR constitutively interacts with the DNA region between *baaR* and *bab2_0216*, repressing divergent transcription on both strands. When present in the environment, adipic acid is likely transported by the MFS MucK transporter Bab2_0213, resulting in derepression of transcription from the BaaR-inhibited promoters. The uptake of structurally related molecules, including ε-aminocaproic acid, ε-caprolactone, and tetradecanedioic acid, also induces transcription, although our data provide evidence that these molecules are transported by a genetically distinct system. Once in the cytoplasm, these molecules are predicted to interact with an unknown metabolic/regulatory process that leads to production of an intracellular ligand that binds to and regulates BaaR. Interaction with a ligand likely induces structural changes in BaaR that result in dissociation from DNA. The subsequent derepression of the *bab2_0213-bab2_0217* locus may increase adipic acid uptake, creating a positive feedback loop. Derepressed expression of the *bab2_0213-bab2_0217* locus also indirectly enhances expression of a paralogous gene set, *bab2_1036-bab2_1046*, while attenuating expression of a potential amino acid ABC transport system (*bab2_0277-bab2_282*).

**Figure 12:**
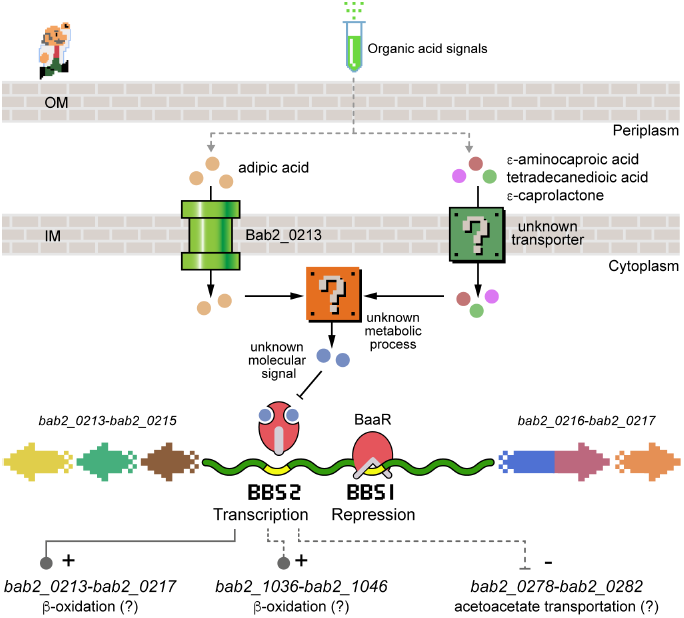
Model of regulation of genes by BaaR based on the results presented in this study. The IclR protein BaaR (Bab2_0215; in red) constitutively interacts with BBS1 and BBS2, repressing transcription of the *bab2_0213-bab2_0216* operon (yellow, green, brown, blue/red, and orange arrows). When present, adipic acid (yellow circles) is specifically transported by the Bab2_0213 transporter (green pipe). The uptake of other molecules, such as ε-aminocaproic acid, ε-caprolactone, or tetradecanedioic acid, may involve an unknown transporter (green (?) box). Once in the cytoplasm, these molecules may activate an unknown metabolic process (orange (?) box), leading to the synthesis of an intracellular molecular signal (blue circles) specifically recognized by BaaR. This interaction is believed to induce structural changes in the BaaR protein and dissociation of the IclR/DNA complex. When transcription of the *bab2_0213-bab2_0217* operon is derepressed, adipic acid uptake is increased and derepression of transcription from the operon is strengthened. This derepression also seems to modulate the *bab2_1036-bab2_1046* and *bab2_0277-bab2_282* locus expression indirectly (gray dashed lines), which in association with *bab2_0213-bab2_0217* may affect the metabolic state of *B. abortus.*

## Materials and methods

All experiments using live *B. abortus* were performed in biosafety level 3 facilities according to U.S. Centers for Disease Control (CDC) select agent regulations at the University of Chicago Howard Taylor Ricketts Laboratory.

### Chromosomal deletion in *B. abortus*

The different *B. abortus* deletion strains (Δ*bab2_0213*, Δ*bab2_0214*, Δ*baaR* [i.e., *bab2_0215*], Δ*bab2_0216*, and Δ*bab2_0217*) were constructed using a double-recombination strategy. Briefly, after PCR amplification using KOD Xtreme Hot start DNA polymerase (EMD Millipore) with *B. abortus* chromosomal DNA as a template, the corresponding PCR products were purified using the GeneJET PCR Purification Kit (Thermo Fisher Scientific). The in-frame deletion alleles (carrying 5’ and 3’ flanking sequences of *B. abortus* locus tags) were digested with restriction enzymes (New England Biolabs) before ligation using T4 DNA ligase (New England Biolabs) or were directly Gibson assembled (New England Biolabs) into the suicide plasmid pNPTS138 (M. R. K. Alley, unpublished data), which carries the *nptI* gene for initial selection and the *sacB* gene for counter-selection on sucrose. Plasmids were then transformed in *E. coli* Top10 strains, and single colonies carrying the plasmid with the correct insert were screened by PCR and Sanger sequenced. Sequenced plasmids were then purified from *E. coli* using the GeneJET Plasmid Miniprep Kit (Thermo Fisher Scientific). The WT *B. abortus* strain 2308 was transformed with pNPTS138 carrying the different deletion alleles by electroporation, and single-crossover integrants were selected on Schaedler Blood Agar (SBA) plates supplemented with kanamycin (50 μg/ml). Counter-selection for the second crossover event was carried out by growing kanamycin-resistant colonies overnight under nonselective conditions and then plating on SBA plates supplemented with 5% sucrose (150 mM). PCR screening to identify colonies containing the deletion alleles was performed, and the products were Sanger sequenced for verification. If necessary, genetic complementation of the deletion strain was carried out by transforming the *B. abortus* deletion strains with the pNPTS138 plasmid carrying the WT allele locus. The primers, restriction enzymes, plasmids, and strains used are listed in Tables S3 and S4.

### RNA extraction and sequencing

Three independent cultures of the *B. abortus* WT and *ΔbaaR* strains (see Table S5 for strain information) were grown overnight in 5 ml *Brucella* broth (BB) (Becton Dickinson, BD) at 37°C with shaking at 220 rpm. The next morning, the OD_600_ of the cultures was adjusted to 0.1 in 3 ml BB and grown for another 5 hours at 37°C with shaking at 220 rpm. Cultures (final OD_600_ ∼0.3-0.4) were then pelleted, and the cells were immediately frozen at −80°C until RNA extraction. A hot-phenol RNA extraction protocol was performed for each pellet. Cell pellets were resuspended in 400 μl commercial phosphate buffered saline (PBS) buffer and transferred to tubes containing 200 μl SDS lysis buffer (20% SDS w/v, 0.5 M EDTA) preheated at 95°C. The tubes were gently inverted and incubated at 95°C for 5 minutes. After incubation, the contents of each tube were transferred to new tubes containing 600 μl acid phenol/chloroform preheated at 65°C, vortexed for 5 seconds, and incubated for 10 minutes at 65°C. Samples were then centrifuged for 10 minutes at 2500 × *g.* After centrifugation, the top aqueous phase was transferred to 600 **u**l acid phenol/chloroform at room temperature, vortexed, and centrifuged for 10 minutes at 2500 × *g.* The top aqueous phase was transferred to 500 μl chloroform/isoamyl alcohol (24:1) at room temperature, mixed, and centrifuged for 10 minutes at 2500 × *g.* The top aqueous phase was then transferred to a new tube containing 500 μl 100% isopropanol. Precipitation of the RNA was performed overnight at −80°C. RNA was then pelleted at maximum speed for 30 minutes at 4°C, and the pellets were washed twice with 70% ethanol, air dried, and resuspended in 80 μl RNase-free water. To remove any co-purified genomic DNA, 10 μl 10X buffer and 10 μl Turbo DNase (Thermo Fisher Scientific) were added to each sample for 2 hours at 37°C. The samples were then purified using purification columns (from the RNeasy MinElute Cleanup kit; Qiagen) and eluted with 30 μl DNase (RNase)-free water. The eluate was further digested on the column by adding 7 μl Turbo DNase, 7 μl 10X buffer, and 56 μl RNase-free water. The reaction was incubated at room temperature for 30 minutes and purified using a Qiagen RNA purification kit according to the manufacturer’s protocol.

For RNA-seq analysis, ribosomal RNA was depleted from WT and Δ*baaR* samples using Ribo-Zero rRNA Removal (Gram-negative bacteria) Kit (Epicentre). Libraries were prepared with Illumina TruSeq RNA kit according to manufacturer’s instructions and were then quantified using a 2100 Bioanalyser (Agilent) and sequenced on a HiSeq2500 (Illumina). The obtained RNA-seq reads were aligned to the genome sequence of *B. abortus* 2308 (RefSeq AM040265) using the readmapper tool in CLC Genomics Workbench (Qiagen) (mismatch cost=2; insertion cost=3, deletion cost=3, length fraction=0.8,similarity fraction=0.8). Differential expression analysis of normalized data and false-discovery rate (FDR) p-values were calculated in CLC Genomic Workbench. RNA-seq data sets are available at the National Center for Biotechnology Information (NCBI) Gene Expression Omnibus (GEO) at accession number GSE107825.

### *lacZ* transcriptional reporter construction

The different *lacZ* reporter genes used in this study were built as follows: the WT DNA fragments corresponding to the different portions of the *bab2_0215 (baaR)-bab2_0216* intergenic region were PCR amplified using *B. abortus* chromosomal DNA as a template. A gBlocks synthetized DNA fragment (Integrated DNA Technology, IDT) correspondingtothe*baaR-bab2_0216* intergenic region carrying mutated BBSs were used as a PCR template to introduce mutations in BBS1 and BBS2. DNA fragments corresponding to *baaR* and *bab2_0216* promoter regions carrying a WT and a mutated BBS were generated by overlapping PCRs. Mutation of BBS2 region 1, region 2 or region 3 were introduced using overlapping primers carrying specific mutations. Primer sequences and information are available in Table S4. As previously described, PCRs were performed using with KOD Xtreme Hot start DNA polymerase (EMD Millipore), gel purified with GeneJET PCR purification Kit (Thermo Fisher Scientific) and digested with restriction enzymes (New England Biolabs). Ligation with the linearized pMR15 plasmid was performed using T4 DNA ligase (New England Biolabs). After ligation and transformation into *E. coli* Top 10 strain, single colonies carrying plasmids with the different inserts were PCR screened and sent for sequencing. The plasmids were then purified with the GeneJET Plasmid Miniprep Kit (Thermo Fisher Scientific) and transformed in WT *B. ovis* or in the different *B. abortus* genetic backgrounds (WT, *Δbab2_0213, Δbab2_0214, ΔbaaR, Δbab2_0216* and *Δbab2_0217)* used in this study. Transformants were then selected on SBA plates supplemented with kanamycin (50 μg/ml). Primer, restriction enzyme, plasmid and strain information are available in Tables S3 and S4.

### Screening for ligands that derepress BaaR in vivo

A WT *B. ovis* strain carrying the pMR15-P_*baaR*_-*lcicZ* reporter plasmid (see Table S5 for strain information) was harvested from a fresh SBA plate supplemented with kanamycin (50 μg/ml) and resuspended at OD_600_= 0.1 in 50 ml of BB containing 50 μg/ml of kanamycin. 100 μl of this *B. ovis* suspension was then used to inoculate each well of the Phenotype MicroArray plates (Biolog). Biolog plates PM1, PM2A, PM3B, PM4A, and PM5 were used for this assay. After growth overnight at 37°C in the presence of 5% CO2, the plates were removed from the incubator, and the OD_600_ and β-galactosidase activities were measured under each growth condition using the Tecan Spark 20M plate reader.

### Additional small molecule screening in *Brucella*

Depending on their solubility characteristics, all ligands used in this study were freshly prepared in DMSO or sterile water. When needed, the pH was adjusted to 7.4. Caproic acid, ε-caprolactam, ε-caprolactone, ε-aminocaproic acid, glutaric acid, adipic acid, pimelic acid, suberic acid, sebacic acid, and tetradecanedioic acid were purchased from Sigma-Aldrich. *Cis,cis* and *trans,trans* muconic acids were purchased from Acros-Organics. Succinic acid was purchased from Fisher-Scientific. Malonic acid was purchased from MP-Biomedicals.

For β-galactosidase transcriptional activity measurements*, B. abortus* liquid cultures were prepared as follows: *B. abortus* strains carrying the different *lacZ* transcriptional fusions were harvested from fresh SBA plates supplemented with kanamycin (50 μg/ml) (strain information is available in Table S5). Cells were resuspended in 1 ml of BB and the corresponding OD_600_ was measured by spectrophotometry (Thermo Fisher Scientific Genesys 20). These *B. abortus* suspensions were used to inoculate (at OD_600_ = 0.1) culture tubes containing 2 ml of BB supplemented with kanamycin (50 μg/ml). These culture tubes also contained different concentrations of small molecules (from 50 μM to 4 mM). After overnight growth at 37°C and 220 rpm shaking, β-galactosidase activity was measured. Each condition was independently tested at least three times using different clones for each time.

### Measurement of β-galactosidase activity

In this study, all β-galactosidase activity measurements were performed in 96-well plates, and the absorbance was measured using the Tecan Spark 20M or Infinite 200 PRO plate reader.

To assess regulation of reporter gene transcription by BaaR, the different *B. abortus* strains were grown on SBA plates supplemented with 50 μg/ml kanamycin. After incubation for 2-3 days at 37°C and 5% CO2, the cells were harvested and resuspended in 1 ml BB; 200 μl of each culture tube were transferred to a clear Corning flat-bottom 96-well plate, and the OD_600_ was measured using the Tecan plate reader. β-galactosidase activities of four different clones for each strain were independently measured at least twice. A representative data set is presented in Figure 6.

For ligand screening on 96-well plates, Phenotype MicroArray (Biolog) plates were prepared as described earlier, and the OD_600_ was measured using the Tecan plate reader. This initial screen was conducted once. For ligand screening in culture tubes, 200 μl each culture tube were transferred to a clear flat-bottom 96-well plate, and the OD_600_ was measured using the Tecan plate reader. In Figures 7 and 8, each condition was independently tested at least three times using different clones for each time.

For β-galactosidase activity measurements, between 2.5 and 10 μl cell suspension were mixed and lysed with 25 μl chloroform in a 96-well chloroform-resistant plate, and 125 μl Z-buffer (60 mM Na2HPO4, 40 mM NaH_2_PO4, 10 mM KCl, adjusted to pH 7) were then added to each well. After addition of 42 μl of a 4 mg/ml O-nitrophenyl-β-D-galactopyranoside solution to each well, the reaction was developed at room temperature and stopped by adding 83 μl of a 1 M sodium carbonate solution. For each reaction, the incubation time was recorded; 200 μl of each reaction were then transferred to a clear flat-bottom 96-well plate, and the OD_42_0 was measured using the Tecan plate reader. Calculation of (β-galactosidase activity was performed using the following formula:

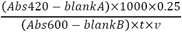

where Abs420 is the absorbance measured at 420 nm of the β-galactosidase reaction, blankA is a blank reaction containing no cells, 0.25 is the total volume of the reaction (ml), Abs600 is the absorbance measured at 600 nm of the cell suspension, blankB is a blank containing *Brucella* broth alone, *t* is the incubation time (min), and *v* is the volume of cells used in the reaction (ml).

### Oxidative stress assay

*B. abortus* WT, *ΔbaaR,* and *ArpoEl* strains (strain information is available in Table S5) grown on SBA plates for 48 hours were harvested and resuspended in Gerhardt’s Minimal Medium, pH 6.8 (GMM) (68). Each sample was then adjusted to a final cell density of 1 **^x^** 10 CFU/ml in 2 ml Gerhardt’s Minimal Medium (GMM), pH 6.8. The test group was subjected to oxidative stress by the addition of 5 mM H2O2 (final concentration); the control group was mock treated with sterile water. After 1 hour of incubation in a shaking incubator at 37°C, the cultures were serially diluted in PBS and plated on tryptic soy agar plates for viable CFU counting. Three independent cultures per condition were prepared for each strain.

### Cell culture and macrophage infection assay

Human monocytic THP-1 cells were cultured in Roswell Park Memorial Institute (RPMI) 1640 medium supplemented with 10% heat-inactivated fetal bovine serum and 2 mM L-glutamine. Differentiation of the cells into inactivated macrophages was induced by the addition of 40 ng/ml phorbol 12-myristate 13-acetate (Sigma-Aldrich) for 48 hours at 37°C in a 5% CO2 atmosphere. Prior to infection, bacteria were harvested from freshly plated SBA plates, resuspended in sterile RPMI medium, and cell densities were adjusted. For infection assays, 5 × 10^4^ THP-1 cells were infected with 5 × 10^6^ cells of the WT or *ΔbaaR B. abortus* strains to achieve a multiplicity of infection (MOI) of 1:100 in 96-well plates. To synchronize the infections after the addition of the *Brucella* cells, the plates were centrifuged at 200 × *g* for 5 min. After 1 hour of incubation at 37°C in a 5% CO2 atmosphere, the medium was removed and replaced with RPMI medium supplemented with gentamicin (50 μg/ml) and incubated for 20 minutes at 37°C in a 5% CO2 atmosphere to kill extracellular bacteria. To determine the numbers of intracellular bacteria at 1, 24 and 48 hours post infection, the cells were washed once with PBS and lysed with 0.1 ml PBS complemented with 0.1% Triton X-100. The lysate was serially diluted and plated on tryptic soy agar plates for CFU counting. This infection experiment was performed in triplicate using two different clones of each strain.

### Protein expression plasmids

DNA fragments corresponding to the full-length BaaR (residues 1-284) protein, the BaaR protein deleted for the first 8 (9-283), 20 (21-284), or 39 (40-284) N-terminal amino acids, and the BaaR LBD (115-284) were amplified by PCR using KOD Xtreme Hot start DNA polymerase (EMD Millipore). The primers used for cloning are listed in Table S4. After purification using the GeneJET PCR Purification Kit (Thermo Fisher Scientific), the PCR products were directly Gibson assembled (New England Biolabs) or digested with restriction enzymes (New England Biolabs) and ligated using T4 DNA ligase (New England Biolabs) into a linearized pET28a plasmid. Fragments corresponding to BaaR (residues 9-283) and BaaR (residues 21-284) were respectively cloned into pMCSG81 and pMCSG73 plasmids using a ligation-independent procedure (69,70). *E. coli* Top10 strains were then transformed with the different plasmids by electroporation, and transformants were selected on Luria Broth (LB, Fisher) agar plates supplemented with 50 μg/ml kanamycin. After PCR screening and sequencing, the corresponding plasmids were purified using the GeneJET Plasmid Miniprep Kit (Thermo Fisher Scientific) and transformed into *E. coli* BL21-Gold(DE3) or Rosetta(DE3)(pLysS) strains (Stratagene) for protein expression (see Table S5 for strain information).

### Protein expression and purification

An overnight LB (Fisher) pre-culture (100 ml) was used to inoculate 1 L LB supplemented withtheappropriateantibiotics. Overexpression of the different His-tagged BaaR proteins was induced at an OD_600_ of ∼0.8 (37°C, 220 rpm) by adding 1 mM isopropylβ-D-1-thiogalactopyranoside (GoldBio). After 5 hours of induction, cells were harvested by centrifugation at 11000 × g for 20 minutes at 4°C. The cell pellets were resuspended in 20 ml of a buffer containing 10 mM Tris-HCl, pH 8.8, 150 mM NaCl and 10 mM imidazole and supplemented with 50 μl of a DNaseI solution at 5 mg/ml and one-half tablet of complete protease inhibitor cocktail (Roche). The cells were then disrupted by one passage in a microfluidizer (Microfluidics LV1). The resulting cell lysate was clarified by centrifugation at 39000 × g for 20 minutes at 4°C. Purification of the His-tagged proteins was performed by nickel affinity chromatography (nitrilotriacetic acid resin; GE Healthcare). After binding of the clarified lysate to the column, three washing steps were performed using 10 mM, 30 mM, and 75 mM imidazole Tris-NaCl buffers, followed by elution with 200 mM and 500 mM imidazole Tris-NaCl buffers. All purification steps were carried out at 4°C. Protein purity was assessed by running the eluate on a 14% SDS-PAGE gel and staining the gel with Coomassie blue. Proteins were then dialyzed overnight at 4°C against 2 L Tris-NaCl buffer (10 mM Tris pH 8.8, 150 mM NaCl, 1.5 mM EDTA). The protein concentrations were estimated using a colorimetric Bradford protein assay method kit (Thermo Fisher Scientific). Protein samples were concentrated using a centrifugal filter (3 kDa molecular mass cutoff, Amicon-Millipore). If necessary, protein aliquots were flash frozen in liquid nitrogen after addition of glycerol to a final concentration of 50%.

### Electrophoretic mobility shift assay (EMSA)

For gel shift assays, fluorescent DNA probes corresponding to different portions of the WT *bab2_0215*-*bab2_0216* intergenic region were PCR amplified using *B. abortus* chromosomal DNA as a template. Upstream primers positioned before BBS1 or BBS2 and a downstream fluorescent primer positioned at the beginning of the *baaR* gene were used to generate the “long” (375 or 289 nucleotides) and “short” (196 nucleotides) fluorescent DNA probes. The fluorescent primer (Integrated DNA Technology, IDT) was labeled with the Alexa Fluor 488 dye with an excitation wavelength of 492 nm and an emission wavelength of 517 nm. Using the same set of primers, a gBlocks DNA Fragment (Integrated DNA Technology, IDT) corresponding to the *bab2_0215-bab2_0216* intergenic region carrying mutated BBSs was also used as a PCR template to introduce mutations in BBS1 and BBS2. The resulting PCR products corresponded to the long (375 nucleotides) or short (196 nucleotides) mutant fluorescent DNA probes. Mutations in BBS2 region 1, 2, or 3 were introduced using three different sets of overlapping primers containing BBS2 region 1, 2, or 3 mutations, respectively. The different fluorescent DNA probes were also used as templates to generate non-fluorescentDNAprobes (short/long/WT/mutated) using a non-fluorescent downstream primer during PCR amplification. Primer sequences and information regarding the gBlocks Gene Fragment are available in Table S5. The PCR products were run on a 1% agarose gel and gel purified using the GeneJET PCR Purification Kit (Thermo Fisher Scientific). DNA concentrations were measured using the Nanodrop One (Thermo Fisher Scientific).

All EMSAs were performed using a BaaR protein corresponding to residues 9-283. Two additional constructs (a full-length (residues 1-284) and a shorter (residues 40-284) BaaR protein) were also used to evaluate the importance of the BaaR N-terminal region. All gel shift assays were performed using previously published protocols (71,72). Briefly, 10 μl reactions were prepared as follows: each reaction consisted of a specific concentration of BaaR protein mixed with 5 ng of a fluorescent DNA probe and 1 μl of a 10× binding buffer (100 mM Tris pH 8.8, 10 mM EDTA, 100 mg bovine serum albumin, 20 mM CaCl_2_, 500 mM KCl, 1 mM dithiothreitol, 50% glycerol, 20 mM MgCl_2_). To control for interaction specificity, fluorescent or non-fluorescentDNAprobes (short/long/WT/mutated) were also added to the mix at defined concentrations. The effect of specific small molecules was evaluated by adding defined concentrations of each molecule to the mix. Protein dialysis buffer was used to bring the total volume to 10 μl. Samples were incubated in the dark at room temperature for at least 30 minutes and then loaded on a fresh 5% native acrylamide gel pre-run for 20 minutes in 1× Tris acetate EDTA buffer (40 mM Tris, 20 mM acetic acid, and 1 mM EDTA). After running the gel for 1 hour at 4°C/110 volts in the dark, the gels were imaged using the BioRad Chemidoc MP imaging system with a 3-minute exposure and the manufacturer’s parameters for Alexa Fluor 488 detection. Each specific condition was tested at least twice.

### Isothermal Titration Calorimetry (ITC) ligand-binding assay

All samples (proteins and ligands) were degassed for 10 minutes prior to ITC measurements, and final dilutions were made using dialysis buffer (10 mM Tris, pH 8.8, 150 mM NaCl, and 1.5 mM EDTA). The ligands were injected into a 200 μl sample cell containing 50 μM purified BaaR LBD (residues 115-284) (protein expression strain information is available in Table S5). A 1 M adipic acid solution was prepared using the dialysis buffer and adjusted to pH 8.8. This same solution was then diluted to 10 mM using the same dialysis buffer. The ligand solution (1 μl) was injected into the cell every 2 minutes, with 20 injections total performed. Measurements were performed twice at 25°C using an iTC200 micro-calorimeter (MicroCal, GE Healthcare).

### Protein expression and purification for crystallization

For crystallization, the *E. coli* strain carrying the pMCSG73 vector was used to overexpress BaaR (residues 21-284) (see strain information listed in Table S5). The pMCSG73 is a bacterial expression vector harboring a tobacco vein mottling virus-cleavable N-terminal NusA tag and a TEV-cleavable N-terminal 6x His and StrepII tag (70). A 2 liter culture of enriched M9 medium was grown at 37°C with shaking at 190 rpm. At OD_600_ ∼1, the culture was cooled to 4°C and supplemented with 90 mg L-seleno-L-methionine (Se-Met, Sigma-Aldrich) and 25 mg of each methionine biosynthetic inhibitory amino acid (L-valine, L-isoleucine, L-leucine, L-lysine, L-threonine, and L-phenylalanine). Protein expression was induced overnight at 18°C using 0.5 mM isopropyl β-D-1-thiogalactopyranoside. After centrifugation, cell pellets were resuspended in 35 ml lysis buffer (500 mM NaCl, 5% (v/v) glycerol, 50 mM HEPES pH 8.0, 20 mM imidazole, and 10 mM β-mercaptoethanol) per liter of culture and treated with lysozyme (1 mg/ml) and 3 ml *E. coli* cells expressing the tobacco vein mottling virus protease. The cell suspension was sonicated, and debris was removed by centrifugation. The Se-Met protein was purified via Ni^2+^-affinity chromatography using the AKTAxpress system (GE Health Systems). The column was washed with 20 mM imidazole (lysis buffer) and eluted in the same buffer containing 250 mM imidazole. Immediately after purification, the His-tag was cleaved at 4°C for 24-48 hours using a recombinant His-tagged TEV protease, resulting in an untagged protein with an N-terminal Ser-Asn-Ala peptide. A second Ni^2+^-affinity chromatography purification was performed to remove the protease, non-cleaved protein, and affinity tag. The purified protein was then dialyzed against 20 mM HEPES pH 8.0, 250 mM NaCl, and 2 mM dithiothreitol buffer. Protein concentrations were determined by UV absorption spectroscopy (280 nm) using the NanoDrop 1000 spectrophotometer (Thermo Fisher Scientific). The purified Se-Met BaaR protein was concentrated to 44 mg/ml using a centrifugal filter (10 kDa molecular weight cutoff, Amicon-Millipore).

### Crystallization

Initial crystallization screening was carried out using the sitting-drop, vapor-diffusion technique in 96-well CrystalQuick plates (Greiner Bio-one). Trays were prepared using a Mosquito robot (TTP LabTech) and commercial crystallization kits (MCSG-1-4, Anatrace). The drops were prepared by mixing equal volumes (0.4 μl) of the purified protein (44 mg/ml) and the crystallization solution equilibrated against 135 μl of the same crystallization solution. After 1 week at 16°C, we obtained monoclinic crystals in condition #95 of the MSCG-4 screen corresponding to 200 mM calcium acetate, 100 mM HEPES pH 7.5, and 10% (w/v) PEG 8000. Prior to flash freezing in liquid nitrogen, crystals were washed for few seconds in the crystallization solution containing up to 12% ethylene glycol for cryoprotection.

### Crystallographic data collection and data processing

Se-Met crystal diffraction was measured at a temperature of 100 K using a 2-second exposure per degree of oscillation. Crystals diffracted to a resolution of 1.95 Å and the corresponding diffraction images were collected on the ADSC Q315r detector with an x-ray wavelength near the selenium edge of 12.66 keV (0.97927 Å) for SAD phasing at the 19-ID beamline (SBC, Advanced Photon Source, Argonne, IL, USA). Diffraction data were processed using the HKL 3000 suite (73). The scaled amplitudes revealed that the crystal belonged to the P2_1_ space group with the following cell dimensions: *a*=74.15 Å, *b*=112.46 Å, *c*=83.65 Å, *β*=115.86^°^ (see Table S1).

The structure was determined by single wavelength anomalous dispersion (SAD) phasing using SHELX C/D/E, mlphare, and dm, and initial automatic protein model building with Buccaneer software, all implemented in the HKL3000 software package (73). The initial model was manually adjusted using COOT (74) and iteratively refined using COOT, PHENIX (75), and/or REFMAC (76); 5% of the total reflections were kept out of the refinement in both REFMAC and PHENIX. The final structure converged to an R_work_ of 17.7% and R_free_ of 21.2% and includes four protein chains (A: residues 21-284, B: 21-283, C: 21-284, D: 20-284) forming two dimers, one ethylene glycol molecule, seven acetate molecules, one calcium ion, and 315 ordered water molecules. The BaaR protein contained three N-terminal residues (Ser-Asn-Ala) that remain from the cleaved tag and were not visible in the structure. The stereochemistry of the structure was checked using PROCHECK (77) and the Ramachandran plot and was validated using the PDB validation server. Coordinates of BaaR have been deposited in the PDB (PDB ID: 5WHM). Crystallographic data and refined model statistics are presented in Table S1. Diffraction images have been uploaded to the SBGrid data server (Data DOI: 10.15785/SBGRID/491).

### Palindromic motifs, protein sequence alignment, and structural homology

Palindromic motifs were identified using the Palindromic Sequence Finder server (http://www.biophp.org/minitools/find_palindromes/demo.php) and the Geneious software. Amino acid sequences were aligned using the M-COFFEE Multiple Sequence Alignment Server (78) and shaded using BoxShade. Figures of the structures, structura alignments, and RMSD calculations were performed using PyMOL (PyMOL Molecular Graphics System, version 1.7.4; Schrödinger, LLC). The XtalPred server (79) and Dali server (80) were used to identify proteins with the highest structural and sequence homologies. A structural model of Bab2_0282 based on the *B*. *malei* PBP structure (PDB ID: 3I09) was generated using the ExPASy SWISS-Model server (81).

## Acknowledgements

We thank the members of the Crosson laboratory for helpful discussions and the members of the SBC at Argonne National Laboratory for their help with data collection at beamline 19-ID. This project was supported by federal funds from NIH-NIAID (grant numbers U19AI107792 and R01AI107159 to SC). JWW was supported by NIH Ruth Kirschstein Postdoctoral Fellowship F32GM109661.

## Author contribution

JH and SC conceptualized and designed the study. JH performed all the experiments, except the RNA-seq experiment, which was performed by DMC and the BaaR crystallization and x-ray structure determination, which was performed by YK and RW. Experiments in BSL3 were performed by JH with technical assistance of DMC, AF and JWW. JH and SC wrote the manuscript. JH, DMC, AF, JWW, YC, GB and SC reviewed and validated the results and approved the final version of the manuscript

## Supplementary files

### Tables

**Table S1:** Data collection and refinement statistics.

**Table s4:** RNA sequencing full data set.

**Table S3:** Phenotype MicroArrays.

**Table S4:** Primers.

**Table S5:** Strains.

### <u>Figures:</u>(see next page)

**Figure S1:**
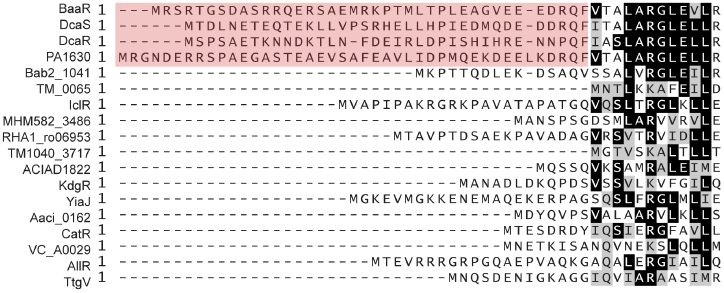
Protein sequence alignment of the N-terminal regions of various IclR proteins with available structure data. Full-length sequences were used for the alignment; only the N-terminal regions are shown. Extra-long N-terminal regions are shaded in light red. The *B. abortus* BaaR amino acid sequence was aligned to *Acinetobacter* sp. ADP1 DcaS (ACIAD1684), *Acinetobacter* sp. ADP1 DcaR (ACIAD1688), *P. aeruginosa* PA1630, *B. abortus* Bab2_1041, *Thermotoga maritime* TM_0065 (PDB ID: 1MKM), *E. coli* IclR (PDB ID: 2O9A), *Microbacterium* sp. HM58-2 MHM582_3486 (PDB ID: 5H1A), *Rhodococcus* sp. RHA1_ro06953 (PDB ID: 2O0Y), Silicibacter sp. TM1040 TM1040_3717 (PDB ID: 3R4K), *Acinetobacter* sp. ADP1 ACIAD1822 (PDB ID: 3D3O), *E. coli* KdgR (PDB ID: 1YSP), *E. coli* YiaJ (PDB ID: 1YSQ), *Alicyclobacillus acidocaldarius* Aaci_0162 (PDB ID: 5TJJ), *Rhodococcus* sp. RHA1 CatR (PDB ID: 2G7U), *Vibrio cholera* VC_A0029 (PDB: 3BJN), *E. coli* AllR (PDB ID: 1TF1) and *P. putida* TtgV (PDB ID: 2XRN).

**Figure S2:**
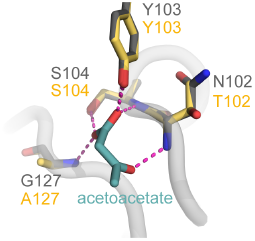
Structural alignment of *B. mallei* PBP (PDB ID: 3I09) to a structural model of Bab2_0282 based on 3I09. Residues (N102, Y103, S104, and G127), involved in the interaction (magenta dashed lines) of the protein with acetoacetate (in green) are shown as gray sticks. The equivalent residues (T102, Y103, S104, and A127) in Bab2_0282 are shown as yellow sticks.

**Figure S3:**
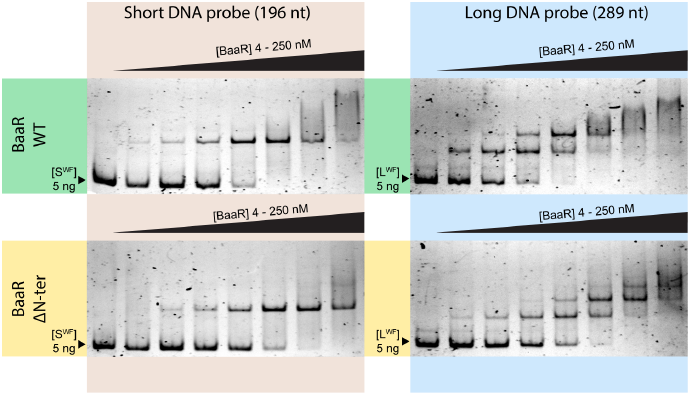
Gel shift assays performed using the WT BaaR (residues 1-284) or BaaR with an N-terminal deletion (Δ1-39). A shift in the size of a short (196 nucleotides long; S^WF^) or long (289 nucleotides long; L^WF^) WT fluorescent DNA probe (5 ng) targeting each of these proteins was assessed on a 5% native acrylamide gel. The protein concentrations used were 0, 4, 8, 16, 31, 62, 125, and 250 nM.

**Figure S4:**
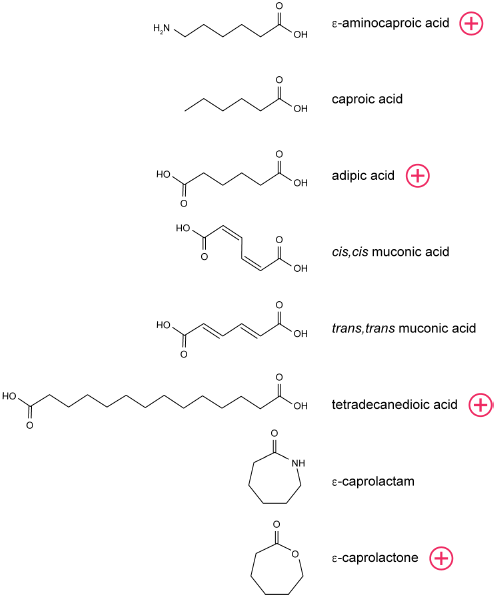
The molecules assessed in this study and their structures. Molecules marked with a red (+) correspond to those affecting the β-galactosidase activity of the *B. abortus* P*_baaR_-lacZ* reporter strain.

